# Precision long-read metagenomics sequencing for food safety by detection and assembly of Shiga toxin-producing *Escherichia coli* in irrigation water

**DOI:** 10.1101/2020.07.17.209718

**Authors:** Meghan Maguire, Julie A. Kase, Dwayne Roberson, Tim Muruvanda, Eric W. Brown, Marc Allard, Steven M. Musser, Narjol González-Escalona

**Author notes:** Corresponding author. Mailing address, Center for Food Safety and Applied Nutrition, Food and Drug Administration, 5001 Campus Drive, College Park, MD 20740, USA.

## Abstract

Shiga toxin-producing *Escherichia coli* (STEC) contamination of agricultural water might be an important factor to recent foodborne illness and outbreaks involving leafy greens. Whole genome sequencing generation of closed bacterial genomes plays an important role in source tracking. We aimed to determine the limits of detection and classification of STECs by qPCR and nanopore sequencing using enriched irrigation water artificially contaminated with *E. coli* O157:H7 (EDL933). We determined the limit of STEC detection by qPCR to be 30 CFU/reaction, which is equivalent to 10^5^ CFU/ml in the enrichment. By using Oxford Nanopore’s EPI2ME WIMP workflow and *de novo* assembly with Flye followed by taxon classification with a k-mer analysis software (Kraken), *E. coli* O157:H7 could be detected at 10^3^ CFU/ml (68 reads) and a complete fragmented *E. coli* O157:H7 metagenome-assembled genome (MAG) was obtained at 10^5^-10^8^ CFU/ml. Using a custom script to extract the *E. coli* reads, a completely closed MAG was obtained at 10^7^-10^8^ CFU/ml and a complete, fragmented MAG was obtained at 10^5^-10^6^ CFU/ml. *In silico* virulence detection for *E. coli* MAGs for 10^5^-10^8^ CFU/ml showed that the virulotype was indistinguishable from the spiked *E. coli* O157:H7 strain. We further identified the bacterial species in the un-spiked enrichment, including antimicrobial resistance genes, which could have important implications to food safety. We propose this workflow could be used for detection and complete genomic characterization of STEC from a complex microbial sample and could be applied to determine the limit of detection and assembly of other foodborne bacterial pathogens.

**IMPORTANCE:** Foodborne illness caused by Shiga toxin-producing *E. coli* (STEC) ranges in severity from diarrhea to hemolytic uremic syndrome and produce-related incidence is increasing. The pervasive nature of *E*. *coli* requires not only detection, but also a complete genome to determine potential pathogenicity based on *stx* and *eae* genes, serotype, and other virulence factors. We have developed a pipeline to determine the limits of nanopore sequencing for STECs in a metagenomic sample. By utilizing the current qPCR in the FDA Bacteriological Analytical Manual (BAM) Chapter 4A, we can quantify the amount of STEC in the enrichment and then sequence and classify the STEC in less than half the time as current protocols that require a single isolate. These methods have wide implications for food safety, including decreased time to STEC identification during outbreaks, characterization of the microbial community, and the potential to use these methods to determine the limits for other foodborne pathogens.

## INTRODUCTION

Shiga toxin-producing *Escherichia coli* (STEC) is a foodborne pathogen capable of causing severe illness, notably hemolytic uremic syndrome (HUS), and death (1–4). STEC-mediated foodborne illness cases and outbreaks are most commonly associated with the O157:H7 serotype; however, non-O157 STEC illnesses are increasingly being reported (5–8). In the United States, O157:H7 is responsible for approximately 95,000 cases per year of which over 2,000 require hospitalization (7). A further 170,000 cases can be attributed to non-O157 STEC exposure. Foodborne transmission accounts for nearly 70% of O157:H7 incidents. Foodborne outbreaks have been increasingly produce-related, from 0.7% in the 1970s to 6% in the 1990s (9). More recently (2004-2013) produce accounts for approximately 18% of foodborne outbreaks and *E. coli* is one of the most common bacterial sources (10–13). Produce can become contaminated during production, packaging, or preparation; however, half of the produce-associated infections are linked to contamination prior to purchase (14). As such, agricultural water, including water used for irrigation, is an important potential source of contamination (https://www.fda.gov/food/outbreaks-foodborne-illness/environmental-assessment-factors-potentially-contributing-contamination-romaine-lettuce-implicated; https://www.fda.gov/food/outbreaks-foodborne-illness/investigation-summary-factors-potentially-contributing-contamination-romaine-lettuce-implicated-fall) (15–20). While prevention and mitigation strategies are beyond the scope of this study, detection of STEC bacteria remains paramount for public health.

A standard and accepted method for the detection of STECs in foods is based on a combined result from qPCR and a traditional microbiological method described in the FDA Bacteriological Analytical Manual (BAM) Chapter 4A (https://www.fda.gov/food/laboratory-methods-food/bam-chapter-4a-diarrheagenic-escherichia-coli). It consists of a 24-hour sample enrichment in modified buffered peptone water with pyruvate at 42°C followed by qPCR detection of the main virulence genes (*stx1* and *stx2*) and the *wzy* gene of the O157 antigen for a total analysis time of 2-3 days. Positive qPCR results undergo further analysis and O157:H7 STEC is confirmed by several rounds of selective plating on tellurite cefixime – sorbitol MacConkey agar (TC-SMAC), chromogenic agar, and trypticase soy agar with 0.6% yeast extract (TSAYE) plates for 2-4 more days of analysis time. Isolates confirmed to be pure cultures are assessed for toxigenic potential again by qPCR. Further analysis by whole genome sequencing (WGS) is used to determine the complete scope of pathogenicity and antimicrobial resistance status which would add 3-5 days. In conclusion, approximately two weeks are needed for STEC confirmation by the BAM method.

WGS is rapidly changing the approach to foodborne illnesses and outbreak investigations (21). WGS is being used to monitor and identify foodborne pathogens (22,23) and the presence of antimicrobial resistance or virulence genes (24,25). Specific information on serotype and pathogenicity as it relates to phylogenic relationships is increasingly important in outbreak scenarios (26,27). Some of the virulence genes detected by WGS mediate attachment and colonization of STECs and can be found in the locus of enterocyte effacement (LEE), including intimin (*eae*) and type 3 secretion system (TTSS) effector proteins (*esp, esc, tir*), non-LEE effectors (*nleA, nleB, nleC*), and other putative virulence genes (*ehxA, etpD, subA, toxB, saa*) (4,25,28,29).

Metagenomics for sample microbial analysis and targeted detection have been used extensively in many sample types (e.g. spinach, chapati flour, and ice cream) (30–33). Analysis is typically either by 16S rDNA profiling or by shotgun metagenomic sequencing (30–35). Many studies have recently started using long read approaches (36–38) for metagenomic studies because it provides finished metagenome-assembled genomes (MAGs) for the most abundant species or bacteria in the microbiome sample (36–38). Closed MAGs provide a better assessment of those genomes and their virulence potential or functionality in that ecosystem. Most of these approaches either used mock communities or did not properly address the limits of the technique for either detection or the initial sample concentration required to get a completely closed or draft genome that would allow for target organism characterization in the sample. Each metagenomic approach provides different levels of analysis. 16S rDNA has the most sensitive detection limit, requiring the lowest initial CFU, but the lowest resolution, reporting at the genus level because some species have almost identical 16S rDNA genes and the fragment of the 16S rDNA used is relatively small (39). Shotgun metagenomic WGS provides a less sensitive detection limit (above 10^3^ CFU/ml), but provides information from species to a strain level, including many functional genes in the microbiome sample analyzed (33,36,40,41). Metagenomic analyses can be made either using short or long sequencing reads technologies. Short-read shotgun metagenomics was most commonly used for microbiome analyses (30–32,35,39,39,42), while the use of long-read metagenomics is on the rise in the last few years (36–38,40) for numerous reasons. These were summarized very concisely by Bertrand et al. (2019) (40), where the authors mention that short-read sequencing presents difficulty in accurately assembling the complex, highly repetitive regions that can range in sizes up to hundreds of kilobases (36), especially when multiple species are present. Classification of these short reads into species bins based on clustering relies on consensus genomes and is not precise enough for strain-level metagenomic assemblies that are necessary for outbreak and traceback scenarios (40).

Nanopore sequencing can produce completely closed genomes, while also offering affordability and portability (24,43). It does not employ a size selection process (unlike those seen with Illumina or Pacific Biosciences sequencers) resulting in longer reads that can help assemble complex genomic areas (e.g. phages and highly repetitive regions). Furthermore, because nanopore sequencing can perform real-time base calling, it allows for semi-real-time analysis when paired with the Oxford Nanopore EPI2ME cloud service that has the “What’s in my pot” (WIMP) workflow (44). WIMP identifies reads by taxa using an algorithm with the Centrifuge software (45) and the RefSeq sequence database at NCBI (https://www.ncbi.nlm.nih.gov/refseq/). It identifies the number of reads matching an organism of interest. These reads can potentially be retrieved from the total reads and analyzed separately. These extracted reads can then be *de novo* assembled and could result in a completely closed MAG for the organism of interest, in our case STEC O157:H7. This approach could dramatically reduce the time to detect and identify an STEC strain in a sample from two weeks to three days from enrichment.

Long-read nanopore sequencing has proven a useful tool to close bacterial genomes in metagenomic samples where the bacterial species are present in approximately equal proportions (approx. 12% or 10^7^ CFU/ml) (36,37). The actual cell numbers in the initial sample needed for successful assembly of the genome of interest has not been determined yet. Nanopore sequencing of mock microbial communities suggests that the technology is capable of detecting as few as 50 cells in the reaction (i.e. 4 reads) (37). This, however, will be not enough reads to make an informative identification of STEC serotype or evaluation of potential risk to human health via presence of important virulence genes. While current protocols utilize WGS after isolating individual colonies by the selective plating methods described, culture-independent detection of O157:H7 STECs in irrigation water (that were FDA BAM-confirmed O157 STEC-positive) by Illumina MiSeq and Oxford Nanopore shotgun sequencing resulted in negative results due to low concentrations of the organism of interest (Gonzalez-Escalona, unpublished results). However, instead of analyzing the water directly, we suggest that using samples after enrichment will have a sufficient STEC concentration to assemble their genome from nanopore sequencing, which will allow for identification of their serotype, virulence composition, and antimicrobial resistance gene (AMR) content (31).

In order to test and empirically determine the limits of the technique, nanopore sequencing for: 1) detection, 2) characterization, and 3) closing genomes of STECs, we artificially contaminated STEC-negative irrigation water with 10-fold dilutions of *E. coli* O157:H7 EDL933. We propose a workflow for detection and quantification (qPCR) for STECs in enriched samples followed by identification and typing by nanopore sequencing. We also developed a script to extract the desired reads by taxa from the total sequenced reads.

## MATERIALS AND METHODS

### Bacterial strains and media

We used a variant of the shiga toxin-producing *E. coli* (STEC) EDL933 O157:H7 strain for all our experiments. This strain was from our collection at CFSAN and is a variant of ATCC 43895 that after several passages in the lab has lost the *stx2* phage. EDL933 was grown in static culture overnight in tryptic soy broth (TSB) at 37°C.

### Preparation of *E. coli* EDL933 inocula for spiking experiments

For artificial contamination, overnight culture (10^9^ CFU/ml) of *E. coli* EDL933 was serially diluted 10-fold in TSB. Dilutions containing approximately 10^9^ CFU/ml through less than 10 CFU/ml were used for spiking studies. Dilutions of the overnight culture spread on tryptic soy agar (TSA) plates were used to calculate the number of CFUs per ml in the original culture.

### Sample processing and artificial contamination

An STEC-negative irrigation water sample (200 ml) from the Southwestern US was enriched by adding an equal volume of 2X modified Buffered Peptone Water with pyruvate (mBPWp) and incubated at 37°C static for 5 hours. Antimicrobial cocktail [Acriflavin-Cefsulodin-Vancomycin (ACV)] was added and incubated at 42°C static overnight (18-24 h), according to Chapter 4A of the BAM. *E. coli* EDL933 dilutions were added in a 1:1 ratio to the enriched irrigation water sample.

### Nucleic acid extraction

DNA from artificially contaminated irrigation water enrichment was extracted by two methods for 1) qPCR and 2) nanopore sequencing. A 1ml fraction of each spiked enrichment sample was processed for qPCR analysis according to the FDA BAM Chapter 4A. Briefly, cells were pelleted by centrifugation at 12,000 x *g* for 3 minutes. The pellet was washed in 0.85% NaCl and resuspended in 1mL sterile water. Samples were boiled at 100°C for 10 minutes then centrifuged to pellet debris. The DNA supernatant was saved. Another 1 ml portion of each spiked enrichment sample was extracted using the Maxwell RSC Cultured Cells DNA kit with a Maxwell RSC Instrument (Promega Corporation, Madison, WI) according to manufacturer’s instructions for Gram-negative bacteria with additional RNase treatment. DNA concentration was determined by Qubit 4 Fluorometer (Invitrogen, Carlsbad, CA) according to manufacturer’s instructions.

### STEC qPCR detection

The presence of STEC EDL933 was determined by qPCR as described in Chapter 4A of the BAM detecting *stx1, stx2*, and *wzy*. Briefly, the DNA supernatants recovered from boiled samples were diluted 1:10 in nuclease-free water and 2μl was added to 28μl master mix containing 0.25μM stx1 and stx2 primers, 0.3μM wzy primers, 0.2μM stx1 probe, 0.15μM stx2 and wzy probes, 1X Internal Positive Control Mix (Cat: 4308323, Applied Biosystems), 1X Express qPCR Supermix Universal Taq (Cat: 11785200, Invitrogen), and ROX passive dye. All primers and probes (Supplementary Table 1) employed in this study were purchased from IDT (Coralville, IA, USA).

### Whole genome sequencing, contigs assembly and annotation

DNA recovered from the *E. coli* EDL933 spiked water enrichment samples was sequenced using a GridION nanopore sequencer (Oxford Nanopore Technologies, Oxford, UK). The sequencing libraries were prepared using the Genomic DNA by Ligation kit (SQK-LSK109) and run in FLO-MIN106 (R9.4.1) flow cells, according to the manufacturer’s instructions for 72 hours (Oxford Nanopore Technologies). The run was live base called using Guppy v3.2.10 included in the MinKNOW v3.6.5 (v19.12.6) software (Oxford Nanopore Technologies).

The initial classification of the reads for each run was done using the “What’s in my pot” (WIMP) workflow contained in the EPI2ME cloud service (Oxford Nanopore Technologies). That workflow allows for taxonomic classification of the reads generated by the GridION sequencing in real time. Using the WIMP classification output, the reads that were identified as *“Escherichia coli*”* were extracted and saved in a single fastq file using a custom python script (v2.7.3) (Supplementary Note 1). The metagenome-assembled genomes (MAGs) for each spiked sample were obtained by *de novo* assembly using 1) all nanopore data output and 2) extracted *E. coli* reads using the Flye program v2.6 (46) with the meta parameter. The assembled contigs were classified by taxonomy by Kraken (47) using GalaxyTrakr (48) (Flye+Kraken). The presence of the complete genome and synteny of the completely closed genomes on the final assemblies was checked using Mauve genome aligner (49).

### Closure of strain EDL933 genome by nanopore sequencing

For bioinformatic quality control purposes we generated a closed genome of the strain used in in the artificial contamination studies. The long-read sequencing library was prepared and run as mentioned above for the spiked experiments. All reads below 5,000 base pairs in length were removed from further analysis. The genome was assembled using Flye v1.6 (46). The nanopore output resulted in 144,000 reads for a total yield of 716 Mb.

### *In silico* serotyping

The major serotype present in each sample was determined by batch screening in Ridom SeqSphere+ v 7.0.6 (Ridom, Münster, Germany) using the genes deposited in the Center for Genomic Epidemiology (https://cge.cbs.dtu.dk/services/) for *E*. *coli* as part of their web-based tool, SerotypeFinder 2.0 (https://cge.cbs.dtu.dk/services/SerotypeFinder/).

### *In silico* identification of virulence genes

The presence of virulence genes was determined by batch screening in Ridom SeqSphere+ software v 7.0.6 (Ridom) using the genes deposited in the NCBI Pathogen Detection Reference Gene Catalog (https://www.ncbi.nlm.nih.gov/pathogens/isolates#/refgene/) and described in Gonzalez-Escalona and Kase (2019) (25).

### *In silico* identification of antimicrobial resistance genes

We identified the antimicrobial resistance genes present in our sequenced genomes using the EPI2ME Fastq Antimicrobial Resistance workflow (Oxford Nanopore Technologies). This workflow consists of three processes, including 1) quality control of the reads, 2) WIMP analysis using centrifuge and NCBI RefSeq database, and 3) detection of antimicrobial genes using the CARD database (50).

### Metagenomic and EDL933 data accession numbers

The metagenomic sequence data from this study and the nanopore data for the EDL933 strain used in this study are available in GenBank under bioproject number PRJNA639799.

## RESULTS

### Determination of the detection limit of the STEC qPCR method

The qPCR detection limit of *E. coli* EDL933 was determined by calculating CFUs per reaction. DNA was extracted from *E. coli* EDL933 culture according to the boil method described in the BAM Chapter 4A for quantification. The starting CFU/ml concentration determined by plating on TSA plates was 1.5 x 10^9^ CFU/ml. Several 10-fold dilutions of the original boil sample were tested in triplicate with the STEC qPCR assay. The *wzy* gene was detected over six orders of magnitude from 30 to 3 x 10^6^ CFU per reaction (correlation coefficient (R^2^) = 0.99 and efficiency (E) = 98%, Figure 1A). Likewise, the *stx1* gene was detected linearly over six orders of magnitude from 30 to 3 x 10^6^ CFU per reaction (R^2^ = 0.99 and E = 96%, Figure 1B). The limit of the STEC qPCR detection, therefore, was 30 CFU per reaction. This means that the minimal number of detectable STEC cells in enrichment culture using this qPCR protocol will be approximately 10^5^ CFU/ml.

**Figure 1.**
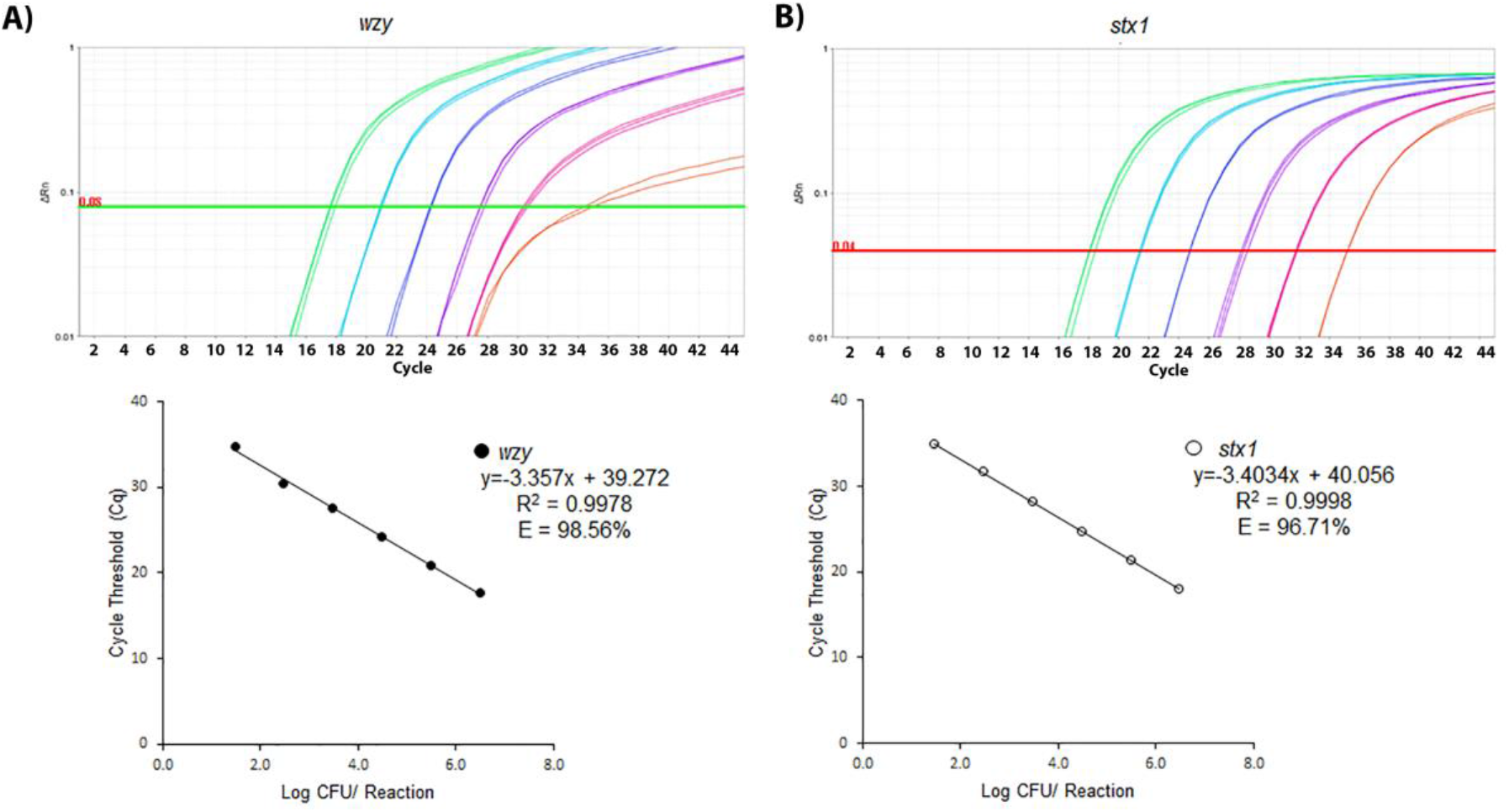
Determination of the detection limit of the qPCR assay. Calibration curves generated using 10-fold dilutions of DNA standards for *E. coli* EDL933 (top) detecting the *wzy* (A) and *stx1* (B) genes. The Cq values were plotted against the log-scale CFU per reaction target concentration (bottom). The R^2^ values and reaction efficiency (E) are also shown.

### STEC-spiked enriched sample preparation for limit of detection of nanopore sequencing

In order to assess the performance and detection limit of nanopore sequencing on STEC enrichment samples, we used an aliquot of irrigation water found by the FDA BAM method to have nondetectable amounts of O157:H7 STEC. Fractions of this enriched irrigation water sample were artificially contaminated with serial dilutions of *E. coli* EDL933 overnight culture. A schematic representation of the workflow is shown in Figure 2. The concentration of the stock *E. coli* EDL933 overnight culture was determined by agar plating to be 1.46 x 10^9^ CFU/ml. Artificial contamination of the enriched field irrigation water sample (1:1), therefore, resulted in an approximate concentration of 7.3 x 10^8^ CFU/ml for the sample referred to as Water+Ecoli1. We confirmed detection and quantification by the BAM qPCR method for this sample. The BAM qPCR method detected the presence of both *wzy* (Cq = 22) and *stx1* (Cq = 23) genes in Water+Ecoli1. Using the standard curve determined above, we calculated 1.1 x 10^5^ CFU/reaction using the *wzy* gene and 9.8 x 10^4^ CFU/reaction using the *stx1* gene. This approximates to 5.4 x 10^8^ CFU/ml using the *wzy* gene and 4.9 x 10^8^ CFU/ml using the *stx1* gene in the Water+Ecoli1 sample. Therefore, the estimated CFU/ml concentration measured by plating and qPCR was very similar.

**Figure 2.**
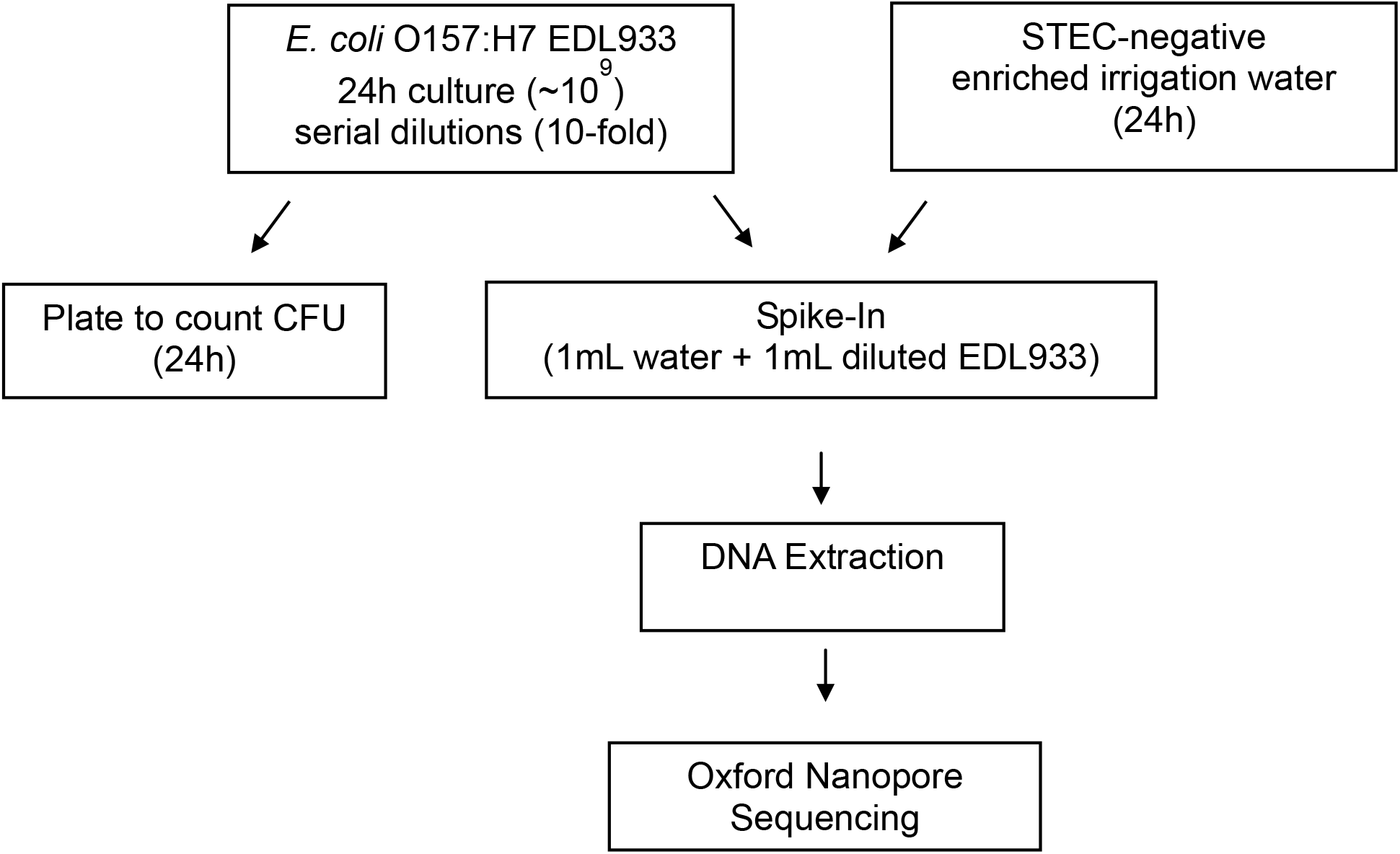
Flow diagram of artificial contamination of enriched, STEC-negative irrigation water through analysis by nanopore sequencing.

### Bacterial community associated with BAM enrichment of irrigation water

The bacterial community composition of the enriched irrigation water was determined by a metagenomic analysis using Oxford Nanopore sequencing of the DNA isolated from the sample. The nanopore output resulted in 5.75M reads in 16.9Gb total yield (Supplementary Table 2). The metagenomic bacterial composition of the sample was analyzed by two methods, Oxford Nanopore EPI2ME “What’s in my pot” (WIMP) workflow analysis and a *de novo* assembly using Flye of all the reads followed by a classification of all the contigs in the assembly by the k-mer software (Kraken). We only reported the taxa accounting for greater than 1% of the total bacterial community. The total WIMP output can be found at https://epi2me.nanoporetech.com/workflow_instance/227465?token=D9EAC8AA-4839-11EA-99BC-71806BDB886C and the total Flye+Kraken output can be found in Supplementary Table 3.

WIMP analysis of the reads obtained by nanopore sequencing of the enriched water sample showed a highly diverse bacterial composition. Even though hundreds of bacterial species were identified in the sample, the majority of the sample was composed of nine bacterial species (>1% in Table 1). The nine bacterial species found in descending cumulative order of frequency were: *Klebsiella pneumoniae* (28.52%), *Enterobacter cloacae* (21.18%), *Enterobacter* sp. ODB01 (6.71%), *Enterobacter kobei* (6.53%), *Pseudomonas putida* (3.88%), *Citrobacter freundii* (3.86%), *Acinetobacter baumannii* (3.69%), *Enterobacter hormaechei* (3.42%), and *Enterobacter xiangfangensis* (1.22%). Additionally, there were 10,806 reads (0.25%) that were identified as belonging to *Escherichia coli*. These *E. coli* reads did not match STEC O157:H7.

**Table 1.** Bacterial community analysis of enriched irrigation water by WIMP. Only species with frequencies above 1% are shown.

|  | Reads | % Reads <sup>a</sup> |
| --- | --- | --- |
| *Klebsiella pneumoniae* | 1,215,922 | 28.52 |
| *Enterobacter cloacae* | 902,950 | 21.18 |
| *Enterobacter sp.* | 286,190 | 6.71 |
| *Enterobacter kobei* | 278,513 | 6.53 |
| *Pseudomonas putida* | 165,323 | 3.88 |
| *Citrobacter freundii* | 164,595 | 3.86 |
| *Acinetobacter baumannii* | 157,421 | 3.69 |
| *Enterobacter hormaechei* | 145,839 | 3.42 |
| *Enterobacter xiangfangensis* | 52,058 | 1.22 |

<sup>a</sup>% Reads were calculated as the proportion of total reads classified by WIMP software analysis

All these microorganisms were identified in the Flye *de novo* assembly. Flye assembly resulted in 677 contigs of different sizes. Taxa identification by Kraken showed that the contigs of bigger sizes belonged to these taxa: *Acinetobacter baumannii* (3,784,399 bp), *Citrobacter freundii* (2,003,414 bp), *Klebsiella pneumoniae* (1,934,072 bp), *Enterobacter cloacae* (1,001,408 bp), *Pseudomonas putida* (438,653 bp), and *Enterobacter kobei* (365,044 bp). *Enterobacter hormaechei* and *Enterobacter xiangfangensis* were also represented in the contigs assembled by Flye, but in the Kraken database *hormachei* and *xiangfangensis* are listed as subspecies of *E. hormachei*. Many other microorganisms were also identified (Supplementary Table 3). Four of the 677 contigs were identified as matching *E. coli* with the largest being 30,837 bp in length.

We were also interested in testing the possibility that AMR genes could be identified. We have used the EPI2ME Fastq Antimicrobial Resistance workflow, which processes all nanopore reads in three stages: 1) reads are passed through a quality filter, 2) taxa are identified by the WIMP workflow, and 3) the classified reads are then analyzed for AMR genes using the CARD database (https://card.mcmaster.ca/home). The prior classification by WIMP permits identification of the AMR genes in each particular species. The AMR genes found in the field irrigation water sample include β-lactamase genes in *Klebsiella pneumoniae* (*bla*_SHV_, *blaA_CT_), Enterobacter cloacae* (*bla*_SHV_, *bla*_ACT_), *Citrobacter freundii* (*bla*_SHV_, *bla*_CMY_), *Acinetobacter baumannii* (*bla*_OXA_), *and Enterobacter hormaechei* (*bla*_ACT_) (https://epi2me.nanoporetech.com/workflow_instance/232304?token=CCD817B6-742C-11EA-B9EA-1AEE73EF14E7). Several efflux pump genes that can be associated with antibiotic resistance were also found in *Klebsiella pneumoniae (acr, ram), Enterobacter cloacae (acr, ram, vga*), and *Acinetobacter baumannii (abe, ade, mex)*. Lastly, a *PhoP* gene mutation was detected in *Klebsiella pneumoniae* (977 reads matching) conferring colistin resistance and the *QnrB23* gene (29 reads matching) that confers fluoroquinolone resistance was detected in *Citrobacter freundii*.

### Nanopore long-read detection limit for *E. coli* spiked into irrigation water enrichment

After establishing the bacterial community of the un-spiked enriched irrigation water sample, we sequenced DNA obtained from the artificially contaminated water enrichment at levels 7 x 10^8^ CFU/ml (Water+Ecoli1) to 7 x 10^3^ CFU/ml (Water+Ecoli6) by nanopore. One microgram of DNA per sample was used as initial DNA amount for nanopore sequencing and each was run in a single flow cell (Figure 2). The total nanopore output per sample can be found in Supplementary Table 2. On average, each run resulted in approximately 5 million reads with a total yield of 17 Gb.

The nanopore output for each sample/run was analyzed using the WIMP workflow and Flye+Kraken as described earlier for the enriched water sample. WIMP workflow classified between 1.7 million (Water+Ecoli1) and 8,500 (Water+Ecoli6) reads as *E. coli* for the serially diluted spiked irrigation water samples (Table 2). These reads accounted for 53 to 0.26% of the total reads for each run (Figure 3). The *de novo* Flye assembly of the reads for each of the spiked water samples produced assemblies with 555 to 780 contigs.

**Figure 3.**
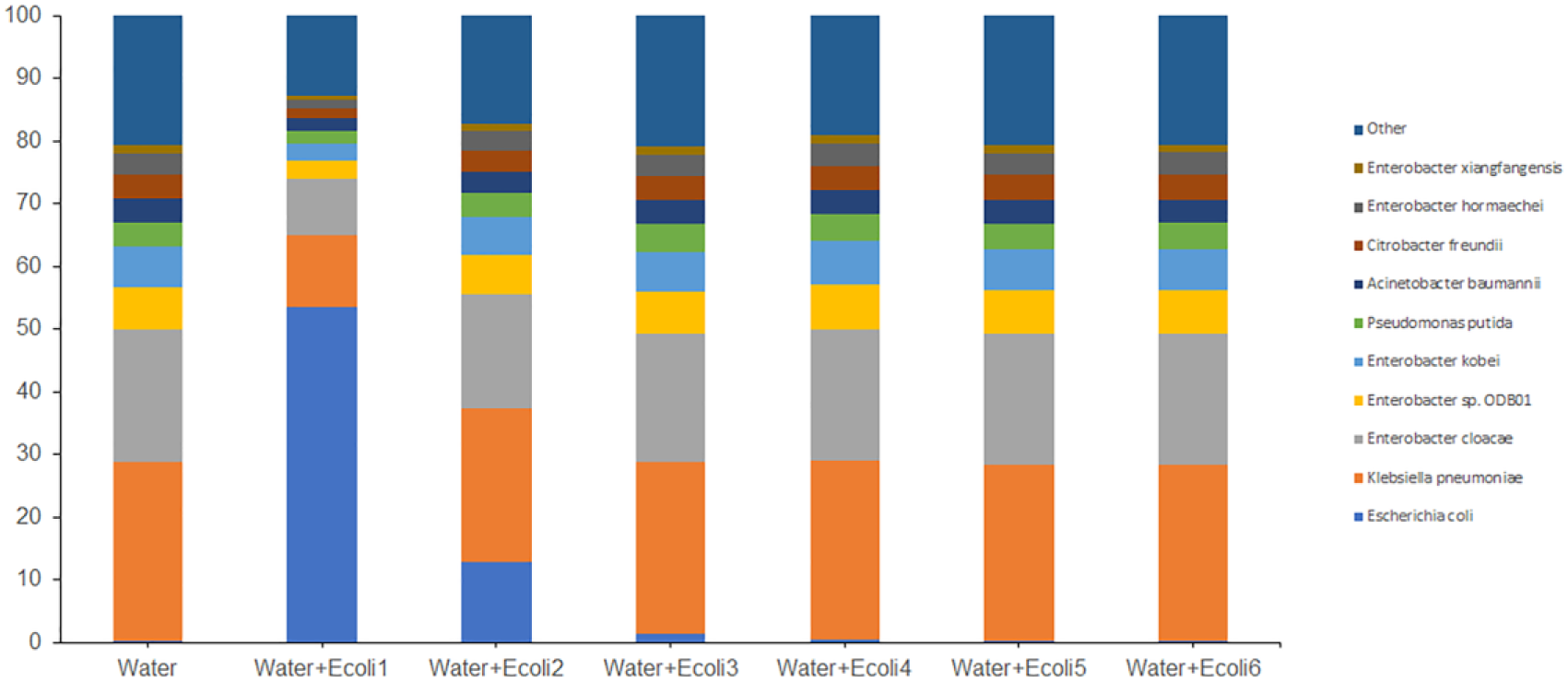
Relative abundance of bacterial species associated with irrigation water un-spiked and spiked with *E. coli* EDL933. Enriched irrigation water (Water) was artificially contaminated with 10-fold dilutions of *E. coli* EDL933 (+Ecoli) with a starting concentration of 7 x 10^8^ CFU/ml (Water+Ecoli1). Reads were analyzed by the EPI2ME WIMP workflow. Bacterial species contributing more than 1% of the classified reads are shown and the sum of the remaining species identified are included as “Other.”

**Table 2.** Virulence of the Flye assemblies obtained with all reads.

| Sample | Innoculation Level (CFU/ml) | Serotype<sup>a</sup> | stx type | eae type | Contig No. (genome and plasmid)<sup>b</sup> |
| --- | --- | --- | --- | --- | --- |
| Water | 0 | O9 | - | - | 677 |
| Water+Ecoli1 | 7.3 x 10<sup>8</sup> | O157:H7 | 1a | gamma-1 | 555 (3+) |
| Water+Ecoli2 | 7.3 x 10<sup>7</sup> | O157:H7 | 1a | gamma-1 | 627 (4+) |
| Water+Ecoli3 | 7.3 x 10<sup>6</sup> | O157:H7/ O9 | 1a | gamma-1 | 753 (35+) |
| Water+Ecoli4<sup>c</sup> | 7.3 x 10<sup>5</sup> | O157:H7/ O9 | 1a | gamma-1 | 780 (80+) |
| Water+Ecoli5 | 7.3 x 10<sup>4</sup> | O9 | - | - |  |
| Water+Ecoli6<sup>d</sup> | 7.3 x 10<sup>3</sup> | O9 | - | - | 644 |

<sup>a</sup>*in silico* serotype using DTU.
<sup>b</sup>In parenthesis, the number of contigs that contain the entire chromosome, a plus sign (+) indicates the presence of a contig that matched the EDL933 circular plasmid.
<sup>c</sup>fragmented genome assembly limit.
<sup>d</sup>*E. coli* O157 detection limit (68 reads matching O157 by WIMP classification).

The enrichment samples with the highest *E. coli* EDL933 spiked concentrations, Water+Ecoli1 and 2, showed the highest number of reads classified as *E. coli* by WIMP (Table 2). As expected, the number of *E. coli* reads classified by WIMP decreased accordingly with dilution of spiked *E*. *coli*. Approximately 8,500 *E. coli* reads were identified in the lowest level of spiked EDL933 Water+Ecoli6 (7 x 10^3^ CFU/ml), almost the same number of reads as the un-spiked enriched water sample (Table 2). WIMP further classified 68 reads as belonging to O157 in that sample. Therefore, the detection limit for O157:H7 by nanopore sequencing was established at 7 x 10^3^ CFU/ml in the enrichment.

### Nanopore long-read genome assembly limit for *E. coli* spiked into irrigation water enrichment

After showing that the detection limit of nanopore sequencing was 7 x 10^3^ CFU/ml, we sought to establish the genome assembly limit. A genome assembly limit is the minimum number of reads used in a *de novo* assembly that produces 1) a complete metagenome-assembled genome or MAG at 20X coverage (chromosome and plasmid, if present) or 2) a fragmented MAG from a complex bacterial background. Obtaining a complete or fragmented *E. coli* O157 MAG will allow us to perform a positive identification of the genome of interest, EDL933, as well as enabling a complete genomic characterization (determine serotype and presence of *stx* types, eae gene, virulence genes, and AMR genes). We expect this read number to be proportionally related to the initial number of CFU/ml in the sample. To test this hypothesis, the completely closed genome of the *E. coli* EDL933 strain used in our spiking experiments was used. By using this reference genome, we ensured the accuracy of our *in silico* analysis to detect the serotype and the entire virulence gene profile. Our EDL933 strain is serotype O157:H7 and has *stx1a, eae* gamma-1 and other virulence genes (*toxB, etpD, tccP*, etc). EDL933 available at GenBank (AE005174) has additionally the *stx*2 gene. We aligned both genomes and found that our version has the entire *stx*2 phage missing (results not shown).

When the total nanopore sequencing output was *de novo* assembled by Flye, the total number of contigs was similar to the un-spiked water sample with 555 and 627 contigs for Water+Ecoli1 and 2, respectively. The *E. coli* EDL933 O157:H7 genome could be detected in 4 or 5 contigs, the plasmid was present as a single contig in each (Table 2). Serotype analysis accurately identified the *E. coli* as O157:H7. The presence of *stx1a* and *eae* gamma-1 were also detected. In fact, the serotype and *stx* and *eae* genes could be determined in Water+Ecoli3 and 4, however the completely closed *E. coli* O157 MAG could not be obtained at any spiked concentration (Table 2).

At the lowest spiked levels in samples Water+Ecoli5 and 6, the number of reads associated with *E. coli* was approximately the same as had been detected in the un-spiked water sample (Table 2). Only the O9 serotype was identified in the assemblies, which was the same as detected in the un-spiked water sample. Likewise, detection of the *stx* and *eae* genes was lost (Table 2). Therefore, we determined that the limit of fragmented assembly was approximately 7 x 10^5^ CFU/ml, but we were not able to obtain a completely O157:H7 STEC closed MAG even at the highest level of 7 x 10^8^ CFU/ml using this approach.

In order to improve the assembly, we decided to extract the *E. coli* reads and perform the *de novo* assembly again. Using WIMP classified reads allowed us to run a script (Supplementary Note 1) that extracted only the reads identified as *E. coli* and perform a similar analysis as above. Flye assembly of these filtered reads produced assemblies that contained fewer number of contigs with larger sizes. High inoculation levels in spiked samples Water+Ecoli1 and 2 resulted in assemblies with 44 and 40 contigs, respectively, each containing the completely closed *E. coli* O157 MAG (chromosome and plasmid each in a single contig) (Table 3). Serotype analysis accurately detected O157:H7 in assemblies from samples Water+Ecoli1 through 4. The same was observed for the recognition of *stx1a* and *eae* gamma-1 genes.

**Table 3.** Virulence of the Flye assemblies obtained with the WIMP *E. coli* extracted reads.

| Sample<sup>a</sup> | WIMP<br>*E. coli* Reads | Serotype<sup>b</sup> | stx type | eae<br>type | Contig No.<br>(genome and<br>plasmid)<sup>c</sup> |
| --- | --- | --- | --- | --- | --- |
| Water | 10,806 | O9 | - | - | 31 |
| Water+Ecoli1 | 1,659,463 | O157:H7 | 1a | gamma-1 | 44 (1+) |
| Water+Ecoli2 | 432,649 | O157:H7 | 1a | gamma-1 | 40 (1+) |
| Water+Ecoli3 | 73,783 | O157:H7/ O9 | 1a | gamma-1 | 41 (8+) |
| Water+Ecoli4<sup>d</sup> | 17,203 | O157:H7/ O9 | 1a | gamma-1 | 92 (63+) |
| Water+Ecoli5 | 10,086 | O9 | - | - | 28 |
| Water+Ecoli6<sup>e</sup> | 8,515 | O9 | - | - | 24 |

<sup>a</sup>CFU/ml levels of EDL933 inoculation can be found in Table 2.
<sup>b</sup>*in silico* serotype using DTU.
<sup>c</sup>In parenthesis, the number of contigs that contain the entire chromosome, a plus sign (+) indicates the presence of a contig that matched the EDL933 circular plasmid.
<sup>d</sup>fragmented genome assembly limit.
<sup>e</sup>*E. coli* O157 detection limit (31 reads matching O157 by Kraken classification).

At the lowest spiked levels in samples Water+Ecoli5 and 6, Flye was able to still produce an assembly but with lower number of contigs (approximately 30 contigs). However, in this case no O157:H7, *stx1a*, or *eae* gamma-1 were detected by *in silico* analyses. Serotype analysis identified the presence of the O9 serotype (Table 3).

Further analysis of the *E. coli* extracted reads for Water+Ecoli5 and Water+Ecoli6 revealed that only 43 and 31 reads matched *E. coli* O157. Kraken identified those reads from a total of 2,326 (Water+Ecoli5) and 1,749 (Water+Ecoli6) *E. coli* reads, respectively. This confirmed our previous analysis that the limit of detection of reads matching O157 is approximately 7 x 10^3^ CFU/ml. Our fragmented assembly limit was still 7 x 10^5^ CFU/ml, but the assembly improved as we were able to produce a completely circular closed chromosome in a single contig with a concentration of at least 7 x 10^7^ CFU/ml.

### Virulence gene identification

In addition to detecting and serotyping STECs, the detection of virulence genes provides necessary information in outbreak scenarios and is important to food safety. We again used the spiked *E. coli* EDL933 genome as reference for *in silico* analysis (Bioproject number PRJNA639799). We have previously reported a set of 94 virulence genes (25) and the EDL933 genome showed the presence of 23 of those genes (Table 4). Among those genes were *esp*, *tccP, nle* genes, *tir*, and *toxB*. The assemblies generated above were analyzed for the presence of all 23 virulence genes. Corresponding to the limits of MAG assembly (either completely closed or fragmented), all virulence genes were detected in Water+Ecoli1 and Water+Ecoli2 when using the total nanopore reads output or extracted reads. On the other hand, we failed to detect *tccP* in Water+Ecoli3 (7 x 10^6^ CFU/ml) and *pssA* and *air* in Water+Ecoli4 (7 x 10^5^ CFU/ml) samples in assemblies with the total nanopore output, but when the *E. coli* reads were extracted from the WIMP output, all virulence genes could be detected. Thus, our extracted *E. coli* reads script improved our genome assembly.

**Table 4.** *In silico* detection of virulence genes in Flye assemblies with all nanopore reads and *E*. coli extracted reads.

|  |  | All reads<br>Water+Ecoli |  |  |  |  |  |  |  | *E. coli* extracted reads<br>Water+Ecoli |  |  |  |  |  |  |
| --- | --- | --- | --- | --- | --- | --- | --- | --- | --- | --- | --- | --- | --- | --- | --- | --- |
| Virulence Gene | EDL933 Reference | Water | 1 | 2 | 3 | 4 | 5 | 6 |  | Water | 1 | 2 | 3 | 4 | 5 | 6 |
| *astA* | + | - | + | + | + | + | - | - |  | - | + | + | + | + | - | - |
| *ehxA* | + | - | + | + | + | + | - | - |  | - | + | + | + | + | - | - |
| *espA* | + | - | + | + | + | + | - | - |  | - | + | + | + | + | - | - |
| *espB* | + | - | + | + | + | + | - | - |  | - | + | + | + | + | - | - |
| *espF* | + | - | + | + | + | + | - | - |  | - | + | + | + | + | - | - |
| *espJ* | + | - | + | + | + | + | - | - |  | - | + | + | + | + | - | - |
| *espK* | + | - | + | + | + | + | - | - |  | - | + | + | + | + | - | - |
| *espP* | + | - | + | + | + | + | - | - |  | - | + | + | + | + | - | - |
| *tccP* | + | - | + | + | - | + | - | - |  | - | + | + | + | + | - | - |
| *etpD* | + | - | + | + | + | + | - | - |  | - | + | + | + | + | - | - |
| *gad* | + | - | + | + | + | + | - | - |  | - | + | + | + | + | - | - |
| *iha* | + | - | + | + | + | + | - | - |  | - | + | + | + | + | - | - |
| *iss* | + | - | + | + | + | + | - | - |  | - | + | + | + | + | - | - |
| *nleA* | + | - | + | + | + | + | - | - |  | - | + | + | + | + | - | - |
| *nleB* | + | - | + | + | + | + | - | - |  | - | + | + | + | + | - | - |
| *nleC* | + | - | + | + | + | + | - | - |  | - | + | + | + | + | - | - |
| *tir* | + | - | + | + | + | + | - | - |  | - | + | + | + | + | - | - |
| *katP* | + | - | + | + | + | + | - | - |  | - | + | + | + | + | - | - |
| *pssA* | + | - | + | + | + | - | - | - |  | - | + | + | + | + | - | - |
| *air* | + | - | + | + | + | - | - | - |  | - | + | + | + | + | - | - |
| *toxB* | + | - | + | + | + | + | - | - |  | - | + | + | + | + | - | - |
| *ecf1* | + | - | + | + | + | + | - | - |  | - | + | + | + | + | - | - |
| IEE | + | - | + | + | + | + | - | - |  | - | + | + | + | + | - | - |

+ Virulence gene detected
- Virulence gene not detected

## DISCUSSION

Considering the importance of irrigation water to food safety, accurate detection and classification of STECs potentially present in those waters is paramount, particularly during an outbreak incident. Current methods of detection include qPCR and extensive selective plating before WGS analysis. This method is a time-consuming process that only provides confirmation of an isolate after almost two weeks of labor. By combining qPCR and long-read metagenomic analysis of the pre-enrichment we can definitively detect an STEC isolate, as well as characterize its virulence potential in 3 - 4 days. While this will not replace eventual confirmation by microbiological methods, this reduces the time for a prospective corrective measure by a complete week.

In our study we have empirically determined the *in silico* limits of detection, classification, and closing genomes of STECs in *E. coli* EDL933 artificially contaminated irrigation water using nanopore sequencing as a proof of concept. We have also developed a pipeline for determination of these limits that can be used for other foodborne or clinical bacteria (Figure 4). Our results showed that the level of STEC O157 needed for detection in the enrichment sample was 10^3^ CFU/ml (Tables 2 and 3). While STEC O157 levels between 10^5^ CFU/ml to 10^6^ CFU/ml were enough to produce a fragmented MAG in a few contigs that allowed for complete characterization of the STEC genome, levels above 10^7^ CFU/ml were enough to produce a completely closed STEC MAG (Table 3) with genome coverage of 385X. The complete plasmid was generated from STEC levels above 10^4^ CFU/ml. These recovered MAGs (either fragmented or completely closed) from all spiked samples above 10^5^ CFU/ml allowed us to comprehensively characterize the virulotype and genome synteny matching 100% to the spiked EDL933 strain (Supplementary Figure 1). The genome of the strain used in this study was sequenced and completely closed by us and used as reference for genome completeness for the *de novo* assemblies sourced from the artificially contaminated samples. Our variant strain of EDL933 was devoid of the *stx*2 phage by qPCR. A comparison of the EDL933 genome published earlier (GenBank accession AE005174) and ours showed that *stx*2 phage was completely missing in our strain, confirming the qPCR results.

**Figure 4.**
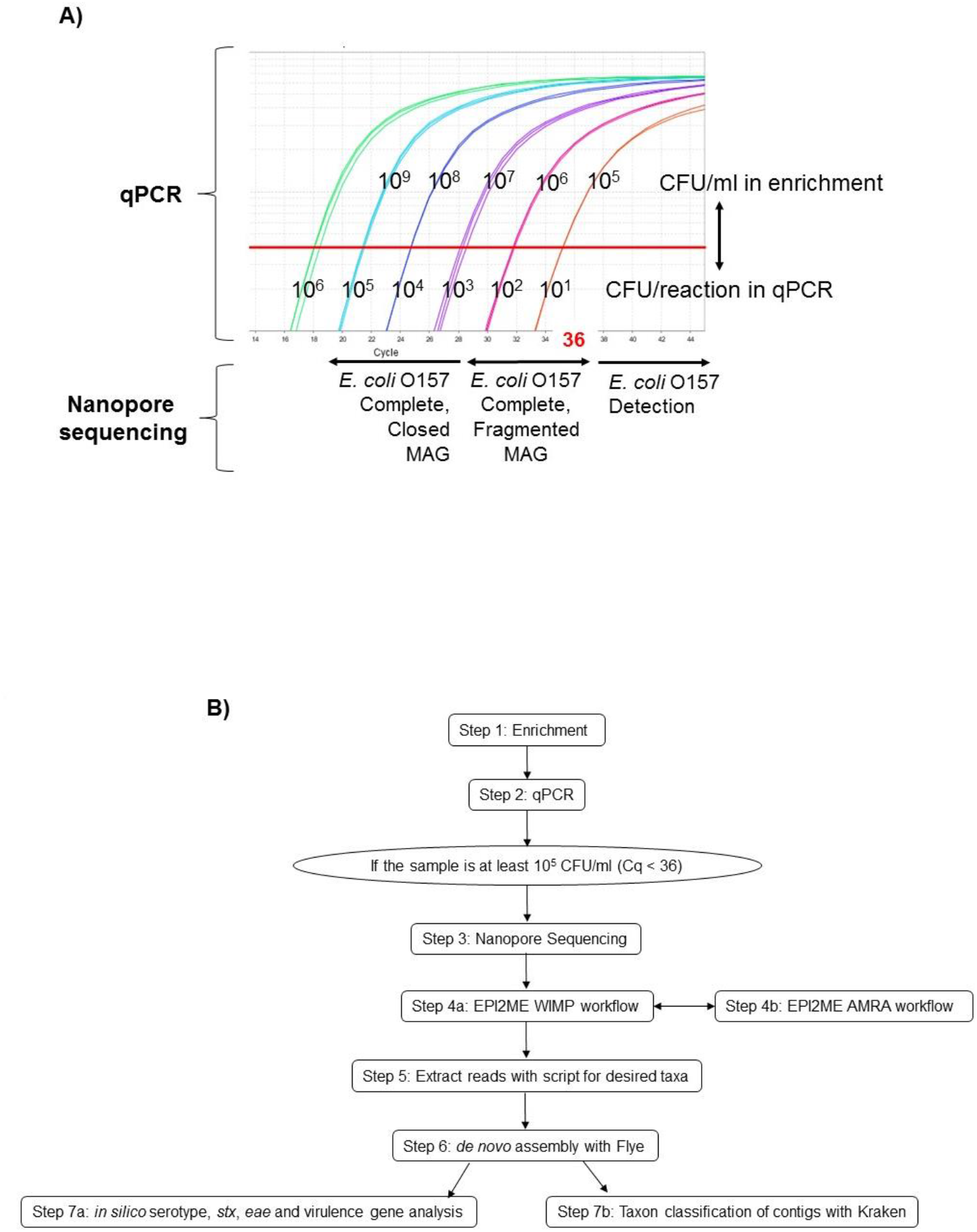
STEC detection and classification by combined qPCR and nanopore sequencing approach. A) Direct comparison of quantitative qPCR detection with *de novo* assembly limits by nanopore sequencing informs detection and classification of STECs. B) Pipeline for detection and classification of STECs in enriched irrigation water using nanopore sequencing and EPI2ME cloud-based services to identify reads of a desired taxa for *de novo* assembly with Flye and *in silico* analysis.

These limits of detection and assembly by nanopore sequencing in conjunction with the use of qPCR for screening the levels of STEC in the pre-enrichments provide an obvious advantage. By combining the qPCR result and the likelihood of genome closure by nanopore sequencing, we have provided an excellent tool for predicting when to pursue sequencing DNA from a particular sample (Figure 4A). The tangible benefit of this combination will depend on the depth of analysis needed – detection versus characterization. While metagenomics has a lower detection limit than that of the BAM qPCR methodology (10^3^ CFU/ml versus 10^5^ CFU/ml) for STEC, it costs 1,000 times more per sample. The sensitivity of nanopore can partially be attributed to the proportion of the initial sample that is used. Nanopore uses approximately 3% of the initial sample (approx. 200 ng/rxn of 6 ug DNA extracted), while qPCR uses 0.02% (2ul of a 1:10 dilution from 1 ml extraction, approx. 1ng/rxn). qPCR provides a directed approach but is only as good as the primers included and can only detect the intended target. In the cases of *Salmonella* spp. or *Listeria monocytogenes*, their detection by qPCR or metagenomic analysis, is beneficial as per FDA’s zero tolerance policy for these two microorganisms in foods. However, presumptive-positive samples would have to be culturally confirmed before any regulatory action could be taken to stop importation or interstate transport of a particular commodity (51) (https://www.fda.gov/media/102633/download; https://www.fda.gov/media/83177/download). On the other hand, if you want to fully characterize an organism by using their complete genome, the methodology described herein will allow any laboratory to expedite the process of detection and characterization (Figure 4B). In the case of STECs the entire genome is needed in order to make an informative decision of their potential health risk to humans. Many *E. coli* strains are not harmful to humans and will pose no risk for public health. Thus, obtaining the completely closed genome of any potential STEC will provide an accurate characterization of all virulence genes it possesses to allow a prediction of potential health risk (25). In our study, the *de novo* assembly of the complete MAG for EDL933 (either in fragments or completely closed) was achieved in all samples with levels above 10^5^ CFU/ml, which was almost equivalent to the limit of detection by qPCR, but with the added benefit of complete genome characterization critically needed during outbreak and traceback investigations.

Mining for specific reads matching your organism of interest in metagenomic sequencing data is challenging and requires conducting assemblies using millions of reads with the consequent time and computing resources that can impact the accuracy of the genome assembler employed (30,31,37,39,43). We took advantage of the WIMP workflow included in the EPI2ME cloud service (Oxford Nanopore) that classifies each single read in a .csv file and downloads those classified reads into a pass folder. A script was written that separated the desired reads by taxa into a new folder. Assemblies produced by using all reads versus using only filtered reads by taxa were compared and completely closed O157:H7 MAGs for 10^7^ and 10^8^ CFU/ml levels with filtered reads were obtained (Tables 2 and 3). As expected, the assemblies produced with taxa filtered reads were faster, more precise and consumed less resources.

Unlike other technologies, nanopore sequencing output is dependent on the quality of the DNA. Some nanopore metagenomics applications can be conducted directly from samples in which DNA extracts do not contain inhibitors and where the target organism(s) is in enough concentration to be detected (36,37). However, STECs in irrigation water require further processing due to low initial concentrations and the presence of considerable humic acid and other unknown inhibitors. Cleaning of those DNAs resulted in loss and shearing of the DNA (Gonzalez-Escalona, unpublished results), with the consequent loss of both resolution and capability of closing target genomes or MAGs. Hence, there is a need to increase the initial biomass of the target organism (STEC in our case) by an enrichment method that will also minimize the contaminants and improve the quality and quantity of the final DNA. By using the Oxford Nanopore Ligation Kit, we have maximized the potential output, which is preferable for metagenomic analyses. In the future we plan to validate these findings with the Rapid Kit, which would further decrease time between sample processing and analysis with the expectation of completely closing MAGs of organisms above 10^7^ CFU/ml. During sample analysis, we noticed that 73% of the reads were below 5000 bp, this also could impact the closing of genomes of interest and also impact the microbial profile or MAGs from that sample. In our case after removing those reads, the same microbial profile was observed, albeit with fewer reads per organism (results not shown). Our future plans include finding a better method for DNA extraction that could provide higher DNA recovery with less shearing to maximize the potential of nanopore sequencing from enriched culture samples. Some authors have addressed DNA shearing when extracting the DNA and suggest gentler bead-beating steps may yield less sheared high molecular weight (HMW) DNA, but might fail in extracting DNA from most difficult organisms (36).

By using our proposed pipeline, we were not only able to improve detection and characterization of our desired organism (STEC), we were also able to identify the bacteria species that were present in the un-spiked enriched irrigation water. This analysis showed that the most common microorganisms (>1% abundance) enriched by the BAM method belonged to the genera *Enterobacter, Klebsiella, Pseudomonas, Acinetobacter*, and *Citrobacter*. We also identified *Salmonella, Escherichia, Serratia, Edwardsiella, Yersinia*, and *Cronobacter* among the 0.1% abundance*. De novo* assembly of the long-read data resulted in 677 contigs with most of these MAGs in a fragmented stage, we were not able to recover a completely closed genome. Nevertheless, we were able to recover the complete genome for *Acinetobacter baumannii* (~3.9 Mb) in 17 contigs (with the longest contig 3,784,399 bp), *Enterobacter cloacae* in 167 contigs, *Klebsiella pneumoniae* in 35 contigs, *Citrobacter freundii* in 21 contigs, and *Pseudomonas putida* in 214 contigs (Supplementary Table 4). We could not recover the MAGs for *Enterobacter sp., Enterobacter kobei, Enterobacter hormaechei subsp. Hormachei* or *Enterobacter hormaechei subsp. Xiangfangensis* strains, even though they were present in higher abundance than *Acinetobacter baumannii*. The most plausible explanation could be that there were many different strains representing those species and therefore it was very hard to assemble their individual genomes. We did find some *E. coli* reads in the un-spiked enriched irrigation water sample (10,806 reads), suggesting the presence of *E. coli* in the original irrigation water sample at very low concentrations. However, the *E. coli* identified by *in silico* molecular serotyping matched to O9 serotype (Table 3), not the O157:H7 serotype of our spiked EDL933 strain, and no virulence genes were found by our *in silico* virulotyping (25). As mentioned by Leonard et al. (2015), having found other non-pathogenic *E. coli* in the original water sample reinforces the need to obtain the complete genomes in order to assess the potential virulence of any *E. coli* strain (31). If we applied the same script for other organisms with abundance above 3% but used different taxa to filter their reads, we could potentially close those genomes as well. This opens up a very attractive way of obtaining closed MAGs from metagenomic samples, similar to what was obtained previously for fecal samples (36).

In addition to taxa identification, another advantage of our proposed pipeline was that we surveyed the microbial community for the presence of AMR genes in the un-spiked enriched water sample. The Antimicrobial Resistance workflow in EPI2ME (Oxford Nanopore) provides AMR gene detection and identifies the organism carrying that AMR gene based on the WIMP classification of the read. This specific result will be hard to achieve when using short reads. AMR genes found in the irrigation water sample included several beta-lactamase and efflux pump genes that confer antibiotic resistance in *Klebsiella pneumoniae, Enterobacter, Citrobacter freundii, Acinetobacter baumannii, and Enterobacter hormaechei*. The *qnrB23* gene variant (29 reads matching) that confers fluoroquinolone resistance was detected in *Citrobacter freundii*. Finally, besides AMR genes we observed the presence of point mutations which confers resistance to colistin in *Klebsiella pneumoniae (PhoP* gene mutation with 977 reads matching, 855X coverage). Antibiotic resistant bacteria in humans has been linked to food sources (52), making the presence of these AMR genes in known human pathogens such as *Klebsiella pneumoniae* and *Acinetobacter baumannii* worrisome. *Acetinobacter* has recently been shown to use killing-enhanced horizontal gene transfer (53), which suggests further study given the high number of AMR genes present in this sample. Additionally, as the soil filters and concentrates the bacteria in the irrigation water, the risk for human consumption increases (54). National and international organizations, such as the National Antimicrobial Resistance Monitoring System (NARMS - https://www.cdc.gov/narms/index.html), One Health approach (https://www.cdc.gov/onehealth/index.html) and the Global AMR Surveillance System (GLASS - https://www.who.int/glass/en/) use the resources of the CDC, USDA, FDA, and WHO to monitor and report the prevalence of and distribute regulatory guidance on antimicrobial resistance in pathogenic and commensal bacteria in food and food animals (52,55). Our pipeline could be an important screening tool to enhance future testing.

## Conclusions

Overall, we tested the limits of detection and assembly for EDL933 O157:H7 in enriched irrigation water using a shotgun long-read sequencing approach. We determined that the limit of detection was 10^3^ CFU/ml. We were able to show that if the levels of the target organism were above 10^5^ CFU/ml, the complete genome will be assembled into several fragments, aided by filtering the reads by taxa. This fragmented genome will be enough to make a complete characterization of the STEC strain including serotype and virulotype. We also determined the detection limit of the BAM STEC qPCR (10^5^ CFU/ml) coincided with our STEC assembly limit by nanopore sequencing. Therefore, we recommend a combination approach using qPCR and nanopore sequencing. In the screening stage, qPCR can provide both detection and an estimate of CFU/ml concentration which could predict if subsequent nanopore sequencing will produce enough data to obtain a complete MAG of the target organism, either closed or fragmented. It also provides information on how many samples can be run in a single flow cell. In our case we used one sample per flow cell, but if levels > 10^7^ CFU/ml we could use 3 - 4 samples per flow cell, reducing the sequencing cost per sample. We expect that the use of this pipeline could enhance the capacity of Public Health entities to respond faster and more accurately during outbreak and traceback investigations. Moreover, further advances in nanopore long-read sequencing accuracy will improve the quality of these MAGs allowing use for phylogenetic analysis and providing a myriad of applications in metagenomics.

## ACKNOWLEDGEMENTS

The study was supported by funding from the MCMi Challenge Grants Program Proposal #2018-646 and the FDA Foods Program Intramural Funds. Dr. Maguire acknowledges a Research Fellowship Program (ORISE) for the Center for Food Safety and Applied Nutrition administered by the Oak Ridge Associated Universities through a contract with the U.S. Food and Drug Administration.

